# High titers of multiple antibody isotypes against the SARS-CoV-2 spike receptor-binding domain and nucleoprotein associate with better neutralization

**DOI:** 10.1101/2020.08.15.252353

**Authors:** Maria G. Noval, Maria E. Kaczmarek, Akiko Koide, Bruno A. Rodriguez-Rodriguez, Ping Louie, Takuya Tada, Takamitsu Hattori, Tatyana Panchenko, Larizbeth A. Romero, Kai Wen Teng, Andrew Bazley, Maren de Vries, Marie I. Samanovic, Jeffrey N. Weiser, Ioannis Aifantis, Joan Cangiarella, Mark J. Mulligan, Ludovic Desvignes, Meike Dittmann, Nathaniel R. Landau, Maria Aguero-Rosenfeld, Shohei Koide, Kenneth A. Stapleford

## Abstract

Understanding antibody responses to SARS-CoV-2 is indispensable for the development of containment measures to overcome the current COVID-19 pandemic. Here, we determine the ability of sera from 101 recovered healthcare workers to neutralize both authentic SARS-CoV-2 and SARS-CoV-2 pseudotyped virus and address their antibody titers against SARS-CoV-2 nucleoprotein and spike receptor-binding domain. Interestingly, the majority of individuals have low neutralization capacity and only 6% of the healthcare workers showed high neutralizing titers against both authentic SARS-CoV-2 virus and the pseudotyped virus. We found the antibody response to SARS-CoV-2 infection generates antigen-specific isotypes as well as a diverse combination of antibody isotypes, with high titers of IgG, IgM and IgA against both antigens correlating with neutralization capacity. Importantly, we found that neutralization correlated with antibody titers as quantified by ELISA. This suggests that an ELISA assay can be used to determine seroneutralization potential. Altogether, our work provides a snapshot of the SARS-CoV-2 neutralizing antibody response in recovered healthcare workers and provides evidence that possessing multiple antibody isotypes may play an important role in SARS-CoV-2 neutralization.

## Introduction

The novel coronavirus, Severe Acute Respiratory Syndrome Coronavirus 2 (SARS-CoV-2), has rapidly spread across the globe, leading to Coronavirus Disease 2019 (COVID-19), devastating mortality, and significant impacts on our way of life. One question that still remains is whether those infected by SARS-CoV-2 generate an immune response that will protect them from reinfection^1^. Moreover, this question is particularly important for the development of a SARS-CoV-2 vaccine, as an effective vaccine would need to generate a potent neutralizing antibody response and immunological memory to provide long-lasting protection^2, 3^. Thus, it is essential that we carefully study and document the neutralizing antibody responses in recovered individuals.

SARS-CoV-2 antibody testing is critical to understanding who has been infected and to provide a picture of seroprevalence in a community^4–9^. However, while these tests are important and provide a relative antibody titer, they are seen more as a “yes or no” type of answer to whether an individual has been infected. Importantly, these tests do not provide information on whether the SARS-CoV-2-specific antibodies present in serum are protective, including through virus neutralization, and as such, a positive antibody test may give individuals a false sense of “immunity” to the virus.

A number of studies have begun to unravel the antibody response to SARS-CoV-2 beyond a simple “yes or no” answer ^9–18^. Here, we hypothesized that by examining the antibody profile in patient’s serum in terms of antigens, antibody isotypes (IgG, IgM, and IgA), and neutralization, we would be able to identify specific signatures associated to effective SARS-CoV-2 neutralization. In this study, we obtained convalescent serum from 101 SARS-CoV-2 PCR-positive healthcare workers and performed a comprehensive analysis of neutralization of authentic SARS-CoV-2 and a pseudotyped lentivirus as well as serum IgG, IgM, and IgA antibody titers to the SARS-CoV-2 spike receptor-binding domain (RBD) and the nucleoprotein (N) (Fig. 1a). Together, this study provides an in-depth look into the SARS-CoV-2 neutralizing antibody response in recovered individuals and highlights the role of multiple antibody isotypes in the development of a potent neutralizing antibody response.

**Figure 1.**
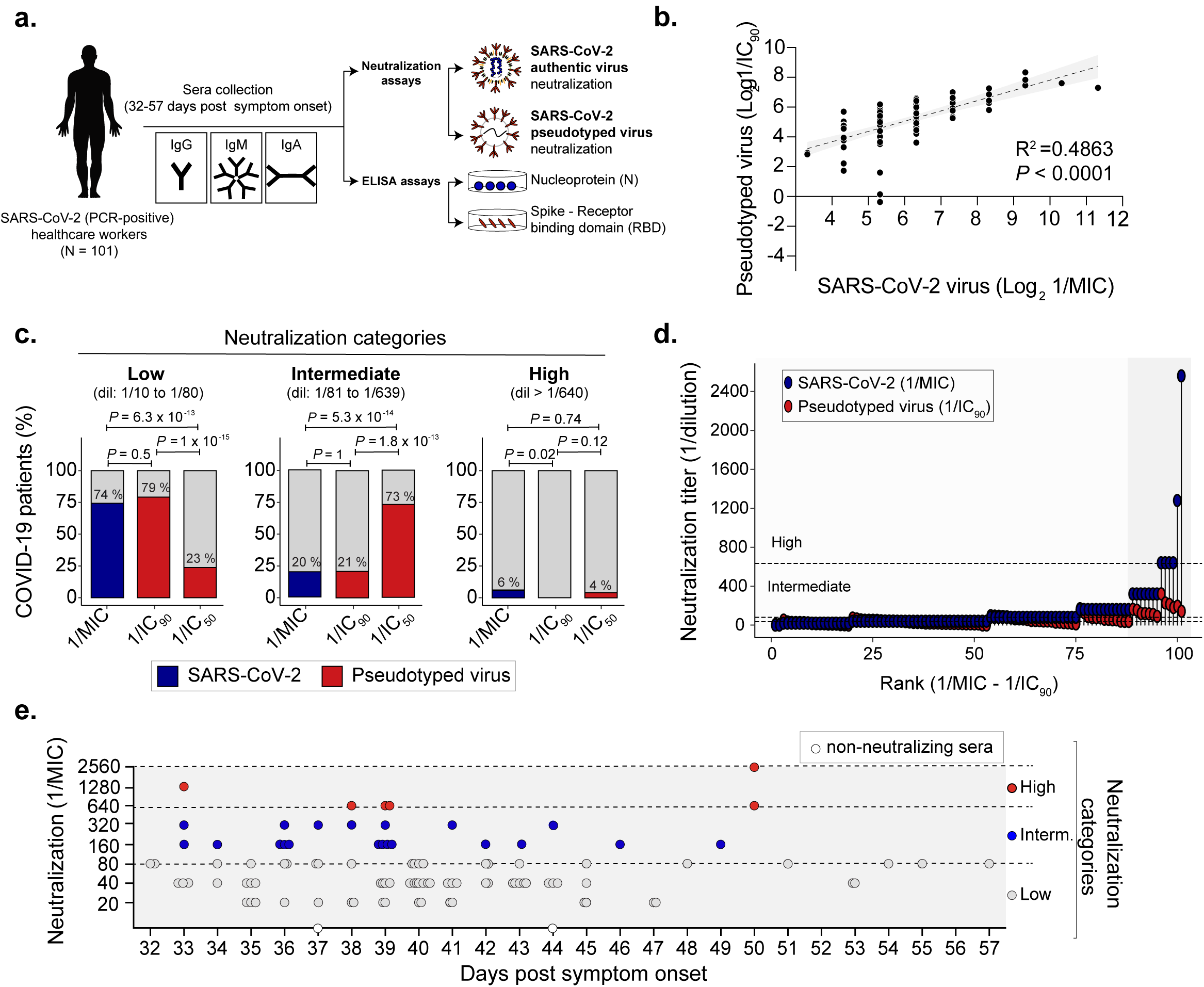
SARS-CoV-2 neutralizing antibody response. **a.** Schematic representation of the experimental desing **b.** Correlation analysis of the sera neutralization level of 101 COVID-19 convalescent patients. The data presented are the log2 of the neutralization titer against the authentic SARS-CoV-2 virus (1/MIC) and the pseudotyped virus (1/IC90). Correlation and linear regression analyses were performed using GraphPad Prism 8. P values were calculated using a two-sided F-test. **c.** SARS-CoV-2 neutralization categories. Sera of convalescent patients was defined as low (dil:1/10 to 1/80), intermediate (dil:1/81 to 1/639) and high (dil > 1/640) based on authentic SARS-CoV-2 virus (blue box) or pseudotyped virus (red box). P values were calculated using two-sided Fisher’s exact test. **d.** Rank of the absolute differences in neutralization capacity from sera of 101 COVID-19 convalescent patients. Neutralization titers against SARS-CoV-2 virus (blue) or pseudotyped virus (red) was ordered based on the absolute difference between 1/MIC and 1/IC90. Dashed lines indicate the different neutralization categories. The gray box highlights those samples with the highest absolute difference in the neutralization capacity between SARS-CoV-2 virus (blue) or pseudotyped virus (red). **e.** Distribution of authentic SARS-CoV-2 virus neutralization titers (1/MIC) over days post symptom onset. Dashed lines indicate the neutralization levels as defined in panel B. White dots indicate two individuals with SARS-CoV-2 non-neutralizing sera.

## Methods

### Serum samples

Serum was collected from 101 healthcare workers from NYU Langone who had laboratory evidence of COVID-19 (PCR-positive) and were at least 21-28 days post symptoms or PCR test. Healthy control sera were collected from patients through the NYU Vaccine Center. All patients gave written consent and all samples were deidentified for this study under IRB #i20-00595 (SARS-CoV-2 infected) and IRB #s18-02037 (healthy controls).

### Plasmids

For pseudotyped virus generation: To construct the SARS-CoV-2 SΔ19 expression vector pcCOV2-Δ19.S, a codon-optimized DNA sequence was synthesized encoding the SARS-CoV-2 Spike (Wuhan-Hu-1/2019). An amplicon was amplified encoding the S protein with a 19 amino acid truncation of the cytoplasmic tail using primers containing flanking 5’-KpnI and 3’-XhoI sites and cloned into pcDNA6 (Invitrogen, Inc.). To construct the ACE2 expression vector pLenti.ACE2-HA, the ACE2 coding sequence was amplified from an ACE2 cDNA (Origene Inc) using primers with flanking 5’-XbaI and 3’-SalI sites and cloned into pLenti.CMV.GFP.puro. For RBD ELISA: The nucleotide sequence of the RBD of SARS-CoV-2 isolate Wuhan-Hu-1 (residues 328-531) was obtained from a GenBank entry MN908947.3. A codon-optimized gene encoding the RBD with a hexa-histidine tag (His6-tag) and biotinylation tag (Avi-tag) at the C terminus was synthesized (Integrated DNA Technologies) and cloned into the pBCAG mammalian expression vector.

### Cells and virus

Vero E6 cells (ATCC CRL-1586) were maintained in Dulbecco’s Modified Eagle’s Medium (DMEM, Corning) supplemented with 10% fetal bovine serum (FBS, Atlanta Biologics) and 1% nonessential amino acids (NEAA, Corning). Human embryonic kidney (HEK) 293T cells were maintained in DMEM supplemented with 10% FBS, and 1% penicillin/streptomycin (P/S). To generate 293T stably expressing ACE2 (ACE2-293T cells), 293T cells were transfected with pLenti.ACE2-HA DNA by lipofection with lipofectamine 2000 (Invitrogen). After 2 days, the cells were selected in DMEM supplemented with 10% FBS, 1% P/S, and 1 µg/ml of puromycin. Single cell clones were expanded and analyzed by flow cytometry for ACE2 expression and a single clone was chosen for subsequent use. Expi293T cells (Thermo Fisher) were maintained in Expi293 Expression Medium (Thermo Fisher). All cells were maintained at 37°C with 5% CO_2_ and confirmed mycoplasma free.

SARS-CoV-2, isolate USA-WA1/2020^19^ (BEI resources # NR52281, a gift from Dr. Mark Mulligan at the NYU Langone Vaccine Center) was amplified once in Vero E6 cells (P1 from the original BEI stock). Briefly, 90-95% confluent T175 flask (Thomas Scientific) of Vero E6 (1×10^7^ cells) was infected with 10 μL of the BEI stock in 3 mL of infection media (DMEM, 2% FBS, 1% NEAA, and 10 mM HEPES, pH 7.0) for 1 hour. After 1 hour, 15 ml of infection media was added to the inoculum and cells were incubated 72 hrs at 37 °C and 5% CO_2_. After 72 hrs, the supernatant was collected and the monolayer was frozen and thawed once. Both supernatant and cellular fractions were combined, centrifuged for 5 min at 1200 rpm, and filtered using a 0.22 μm Steriflip (Millipore). Viral titers were determined by plaque assay in Vero E6 cells. In brief, 220,000 Vero E6 cells/well were seeded in a 24 well plate, 24 hrs before infection. Ten-fold dilutions of the virus in DMEM were added to the Vero E6 monolayers for 1 hour at 37 °C. Following incubation, cells were overlaid with 0.8% agarose in DMEM containing 2% FBS and incubated at 37 °C for 72 hrs. The cells were fixed with 10% formalin, the agarose plug removed, and plaques visualized by crystal violet staining. All experiments with authentic SARS-CoV-2 were conducted in the NYU Grossman School of Medicine ABSL3 facility.

### Pseudotyped virus preparation

Pseudotyped virus was produced by calcium phosphate cotransfection of 293T cells with pMDL, plenti.GFP-NLuc, pcSARS-CoV-2-SΔ19 and pcRev at a ratio of 4:3:4:1. The supernatant was harvested 2 days post-transfection, passed through an 0.22 µm filter and then pelleted by ultracentrifugation for 90 min at 30,000 rpm in an SW40.1 rotor. The pellet was resuspended in 1/10^th^ the original volume of DMEM supplemented with 10% FBS and frozen at −80°C in aliquots.

### SARS-CoV-2 neutralization assay

Vero E6 cells (30,400 cells/well) were seeded in a 96 well plate 24 hrs before infection so that a monolayer was present the following day. Serum samples from COVID-19 convalescent individuals and healthy donors were two-fold serially diluted (spanning from 1:10 to 1:10,240) in DMEM (Corning), 1% NEAA (Corning) and 10 mM HEPES (Gibco). Diluted serum samples were mixed 1:1 (vol/vol) with SARS-CoV-2 virus (6.8 × 10^3^ PFU/ml), and incubated 1 h at 37 °C. During the incubation period, Vero E6 monolayers were washed once with DMEM (Corning) to remove any serum present in the media that could interfere with the assay. After incubation, 100 μl of the serum:SARS-CoV-2 mixtures were added to the Vero E6 monolayers, and cells were incubated at 37°C. Cells were monitored every day for cytopathic effects (CPE) induced by viral infection, and at 5 days post infection cells were fixed in 10% formalin solution (Fisher Scientific) for 1 hr. Cells were then stained by adding 50 µl crystal violet/well and incubated for 30 min. Each well was scored as “0” if there was no monolayer left, “0.5” if there was some monolayer and the well was clearly infected, or “1” if the monolayer was intact. The minimum inhibitory concentration (MIC) was defined as the minimal serum dilution in which the cell monolayer was intact (score = 1). Each serum sample was measured in technical duplicates.

### Pseudotype virus neutralization assay

To determine neutralizing serum titers, ACE2-293T cells were plated in 96 well tissue culture dishes at 10,000 cells/well. The following day, 2-fold dilutions of the donor sera were made in culture medium spanning a range from 1:10 to 1:10240. Each dilution (50 μl) was mixed with 5 μl SARS-CoV-2 SΔ19 lentiviral pseudotype. The mixtures were incubated for 30 min at room temperature and then added to the plated ACE-2 293T cells. The plates were cultured for two days at 37°C with 5% CO2 after which the supernatant was removed and replaced with 50 μl Nano-Glo Luciferase Substrate (Promega, Inc.). Light emission was measured in an Envision 2103 Multi-label plate reader (PerkinElmer, Inc.).

### SARS-CoV-2 Nucleoprotein ELISA

Serum IgG, IgA, and IgM antibodies to the SARS-CoV-2 nucleoprotein were tested using the SARS-CoV-2 IgG, IgA and IgM ELISA kits manufactured by Virotech Diagnostics GmbH for Gold Standard Diagnostics (Davis, CA) following the manufacturer’s instructions. For the detection of IgG and IgA, serum samples were diluted 1:100 in dilution buffer and for IgM, serum samples were diluted 1:101 in RF-Adsorption dilution buffer mixture and incubated at room temperature for 15 min before being added to the wells. Results are reported qualitatively as negative (<9.0 units), equivocal (9.0-11.0 units), and positive (>11.0 units).

### SARS-CoV-2 spike receptor-binding domain (RBD) purification and ELISA

The Expi293F cells (Thermo Fisher) were transiently transfected with the expression vector using the ExpiFectamine™ 293 Transfection Kit (Thermo Fisher, A14524)) and the Expi293 Expression Medium (Thermo Fisher, A14351). The transfected cells were cultured at 37 °C with 8% CO_2_ for 7 days. The culture supernatant was harvested by centrifugation, supplemented with protease inhibitors and clarified by further centrifugation at 8,000 rpm for 20 min and filtration through a 0.22 µm filter. The supernatants were dialyzed into 20 mM sodium phosphate pH 7.4 with 500 mM sodium chloride and the recombinant RBD was purified by immobilized metal ion affinity chromatography (IMAC) using a HisTrap excel column (GE Healthcare). The purified protein was biotinylated using the *E. coli* BirA enzyme produced in house in the presence of 10 mM ATP and 0.5 mM biotin. The RBD protein was purified by IMAC and dialyzed against PBS and stored at − 80 °C. High purity of the purified protein was confirmed using SDS-PAGE, and analysis by size exclusion chromatography using a Superdex 75 10/300 Increase column (GE Healthcare), which showed a single, monodisperse peak consistent with its molecular mass (Extended Data Fig. 1A and B).

For the ELISA, the wells of 384-well ELISA plates (NUNC Maxisort cat# 464718) were coated with 15 µl of 4 µg/ml neutravidin (ThermoFisher cat # 31000) for one hour at room temperature (R.T.) in a humidified chamber. The wells were washed with 100 µl PBST (PBS containing 0.1 % Tween 20) (ThermoFisher, cat# BP337-500) three times using a BioTek 405TS plate washer housed in a BSL-2 biosafety cabinet. The wells were blocked with 0.5 % BSA (Gemini Bio cat# 700-100P, skim milk was not used for blocking because milk can contain biotin that would inhibit antigen immobilization) in PBS overnight at 4 °C. After removing the blocking buffer, 15 µl of 20 nM RBD-His6-Avi-biotin in PBS was added to each well using a Mantis dispenser (Formulatrix). The plates were incubated at R.T. in a humidified chamber for 20 min, and the wells were washed with 100 µl PBST three times using the plate washer. The wells were further blocked with 15 µl of 10 µM biotin in 3% skim milk in PBST (Sigma, cat# 1.15363.0500) for ten min at R.T. in order to block unsaturated neutravidin.

Serum samples were heat-treated at 56°C for one hour^9^ and diluted 158-fold in 1% skim milk in PBST and placed in a 96-well polypropylene plate (ThermoFisher, cat # AB-0796), which served as a master plate. The following sample handling was performed using a Hudson SOLO liquid handler housed in a BSL-2 biosafety cabinet. Twelve microliters of diluted serum were transferred per well of an RBD-immobilized 384-well plate after removing the biotin solution. To prepare 500-fold diluted samples, 8.22 µl of the diluent (1 % skim milk in PBST) was first dispensed into a well to which 3.78 µl of diluted serum from the master plate was added. After 2 hours of incubation at R.T., the plate was washed with PBST three times using the plate washer. 15 µl of secondary antibody (anti human Fab-HRP: Jackson ImmunoResearch, cat# 109-035-097; anti human IgG-HRP: Sigma, cat# A6029-1ML; anti human IgM-HRP: Jackson ImmunoResearch, cat# 109-035-129; and anti-human IgA-HRP: Jackson ImmunoResearch, cat# 109-035-011; x1/5000 diluted in 1 % milk in PBST) was added using the Mantis dispenser and incubated for one hour at R.T. After washing the plate with PBST three times and then with PBS three times, 25 µl of the substrate (SIGMAFAST™ OPD; Sigma P9187-50SET) was added into the wells using the Mantis dispenser. Subsequently, the stop solution (25 µl of 2 M HCl) was added using the Mantis dispenser. The dispenser was programmed in such a way that the reaction in each well was stopped after 10 min. Absorbance at 490 nm was measured using a BioTek Epoch 2 plate reader. All samples were analyzed two independent times to identify outliers. Those yielding inconsistent results in both assays were excluded from analysis (Sample 76 and 86, Extended Data Table 1). Thresholds were determined comparing the 101 SARS-CoV-2 PCR positive samples at two dilutions 1/158 and 1/500 and compared with 43 SARS-CoV-2 PCR negative individuals. The cutoff values were defined as the mean plus three times the standard deviation (SD) of the negative control samples. Results were reported as positive if the values are > 1.091 for IgG, > 0.256 for IgA and > 0.694 for IgM (Extended Data Fig. 1C).

### Data analysis and statistics

All experiments were performed in technical duplicates and data analysis and statistics were performed using GraphPad Prism (Version 8.4.3), R Studio (Version 1.2.5001), and R (Version 3.6.3).

### Data availability

Data that supports all figures, extended data, and analysis is found in Extended Data Table 1.

## Results

### SARS-CoV-2 seroneutralization capacity is low in the majority of individuals

To better understand the human SARS-CoV-2 neutralizing antibody response in convalescent patients, we obtained serum from 101 COVID-19-recovered New York City healthcare workers who had experienced symptoms and had tested positive (by PCR testing) for SARS-CoV-2 in March 2020 (Fig. 1a and Extended Data Table 1). To begin, we assessed how well each individual’s serum was able to neutralize both authentic SARS-CoV-2 (isolate USA WA01/2020), and a lentiviral pseudotyped virus bearing the SARS-CoV-2 Spike protein (pseudotyped virus) *in vitro*. Pseudotyped viruses are a safe alternative to authentic virus assays20, and can therefore be employed by a greater number of research institutes and clinical laboratories to assess neutralization of convalescent serum. However, several studies have shown differences in sensitivity between neutralization of authentic SARS-CoV-2 and specific pseudotyped viruses^21, 22^. Thus, we set out to determine serum neutralization capacity using both our in-house lentiviral pseudotyped virus and the WA01/2020 strain of SARS-CoV-2.

For the neutralization assays performed with authentic SARS-CoV-2 virus we determined the minimum inhibitory concentration (MIC), calculated as the minimum serum dilution at which serum is fully-protective (Extended Data Table 1). For the neutralization experiments performed with the pseudotyped virus, we used a reporter pseudotyped virus expressing luciferase which allows for the calculation of the IC90 and the midpoint transition (IC_50_) (Extended Data Fig. 2 and Extended Data Table 1). Using these systems, we found that neutralization of the authentic SARS-CoV-2 and the pseudotyped virus was positively correlated (Fig. 1b, R^2^ = 0.49, p<0.0001). Moreover, the SARS-CoV-2 neutralizing antibody response could be broken into three categories (Fig. 1c). Using 1/MIC and 1/IC_90_ we find the majority of infected individuals (~75%) had a low neutralization capacity, roughly 20% of individuals had intermediate neutralization and only a select few (~5-6%) had high neutralization of the authentic virus. When comparing pseudotyped virus to authentic SARS-CoV-2 virus neutralization, there was no significant difference in the proportion of individuals assigned as low or intermediate (Fig. 1c). In contrast, there is a significant difference between the percentage of individuals deemed high neutralizers using authentic SARS-CoV-2 versus pseudotyped virus (Fig. 1c and d, p=0.02). These results validate the use of the pseudotyped virus as an efficient alternative to determine the neutralizing potential of serum. But they also suggests that authentic SARS-CoV-2 virus may be better able to detect potent neutralizing serums, which has implications for the selection of donors for passive immunization therapy.

Finally, because samples were collected from individuals at varying times post onset of symptoms, it may be expected that the proportion of highly or moderately neutralizing sera would decrease over time. However, we found that there was no particular pattern in this cohort. Individuals with low, medium, and high neutralizing sera determined using authentic SARS-CoV-2 are found between 32 and 57 days (Fig. 1e). Together, these results show that at the time points we analyzed (32-57 days post symptom onset), serum SARS-CoV-2 neutralizing antibody capacity is low in most recovered individuals.

### Human SARS-CoV-2 infection generates antigen-specific, multi-isotype antibody response

Given the broad neutralization capacity observed in this cohort, we were interested in examining the antibody profile of each individual. It is not uncommon that antibody responses are skewed toward one or a few viral proteins^10, 23^. If this is the case, using a single antigen may result in a biased test that inadequately detects antibodies produced by convalescent individuals. Thus, we quantified serum antibody IgG, IgA, and IgM titers by ELISA, focusing on antibodies generated to two SARS-CoV-2 antigens (Extended Data Table 1). First, we used the Gold Standard ELISA assay currently deployed at the Tisch Hospital Clinical Labs in New York City which uses the nucleoprotein (N). Second, we developed an in-house ELISA for the SARS-CoV-2 Spike receptor-binding domain (RBD) (Extended Data Fig. 1). If antibody titers against Spike RBD and N correlate, this suggests that individuals mount a uniform response to both these antigens. A lack of correlation suggests that individuals mount a skewed response toward one of these antigens preferentially.

When we compared the isotype responses to anti-RBD and anti-N directly, we found that IgG correlated the strongest (R^2^ = 0.51) followed by IgA (R^2^ = 0.23) and IgM (R^2^ = 0.21) (Fig. 2a–c, left panels). The percentage of IgG and IgM positive individuals, as detected by reactivity against RBD or N, was similar (Fig. 2a and b, right panels). Strikingly, the majority of individuals with positive titers of IgA to the Spike RBD were largely IgA-negative for N (Fig. 2c, left panel). This translated to a significant difference in IgA detection between the Gold Standard test and our in-house ELISA (Fig. 2c, right panel, p = 2 × 10^-16^), and suggests that the IgA response is largely skewed towards the Spike RBD in comparison to N (Fig. 2c). It is possible that days post symptom onset plays a role in both serum antibody titers and the relative percentage of IgG, IgM and IgA positive people. Within our cohort we observed no relationship between time post symptom onset and antigen-specific antibody isotype titers (Extended Data Fig. 3), even finding IgM present at 50 days post infection (Extended Data Fig. 3B). Together these results suggest that the antibody isotype response to SARS-CoV-2 may be antigen-specific, with IgA skewed towards the RBD.

**Figure 2.**
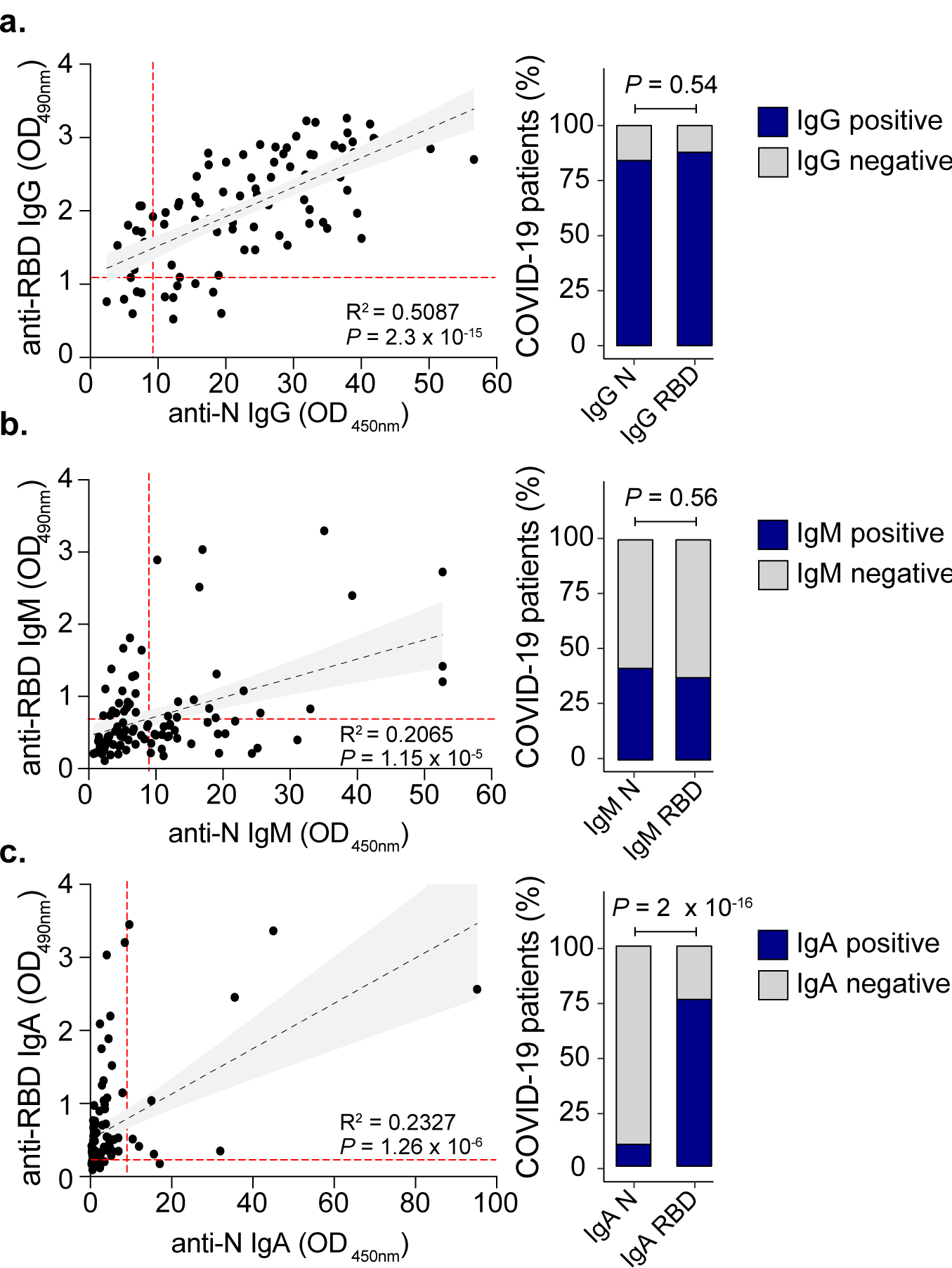
Isotype composition of SARS-CoV2 convalescent serum. **Anti-RBD and anti-N ELISA correlation for IgG (a)**, IgM **(b)** and IgA **(c)**. Left panels: Correlations and linear regressions comparing anti-RBD and anti-N for each antibody isotype (N=101). Analyses were performed using GraphPad Prism 8. P values were calculated using a two-sided F-test. The red dashed lines indicate the threshold (anti-N ELISA for -IgG, -IgM or -IgA are OD450nm = 9; anti-RBD ELISA for -IgG is OD490nm = 1.091; -IgM is OD490nm = 0.256 and -IgA is OD490nm = 0.694) for each ELISA. Right panels indicate the percentage of COVID-19 PCR-positive patients that are positive (blue bars) or negative (gray bars) for IgG **(a)**, IgM **(b)** or IgA **(c)**. P values were calculated using two-sided Fisher’s exact test.

### SARS-CoV-2 neutralization best correlates with RBD antibody response

To better understand how each antigen-specific antibody isotype correlated with neutralization, we compared the ELISA assay antibody titers with virus neutralization. While all anti-RBD isotype titers correlated similarly with the neutralization level of the authentic SARS-CoV-2 (R^2^ = 0.4786 – 0.5194) (Fig. 3a), we found that pseudotyped virus neutralization correlated best with IgG titers (R^2^ = 0.4561) compared with IgA (R^2^ = 0.2556) or IgM (R^2^ = 0.2391) (Fig. 3b). One explanation for these findings may be that there are molecular differences in folding between the RBD of the authentic virus, pseudotyped virus, and purified RBD such that they are recognized differently by antibody isotypes in these assays^24, 25^. Interestingly, with the exception of anti-N IgG antibodies for the authentic virus (R^2^ = 0.5277; R^2^ = 0.255), other anti-N isotype titers showed weaker correlations for either the authentic SARS-CoV-2 virus or the pseudotyped virus (Extended Data Fig. 4). These data suggest that ELISA methods based on the RBD may benefit from detection of additional isotypes, rendering them better suited as predictors of sera neutralization.

**Figure 3.**
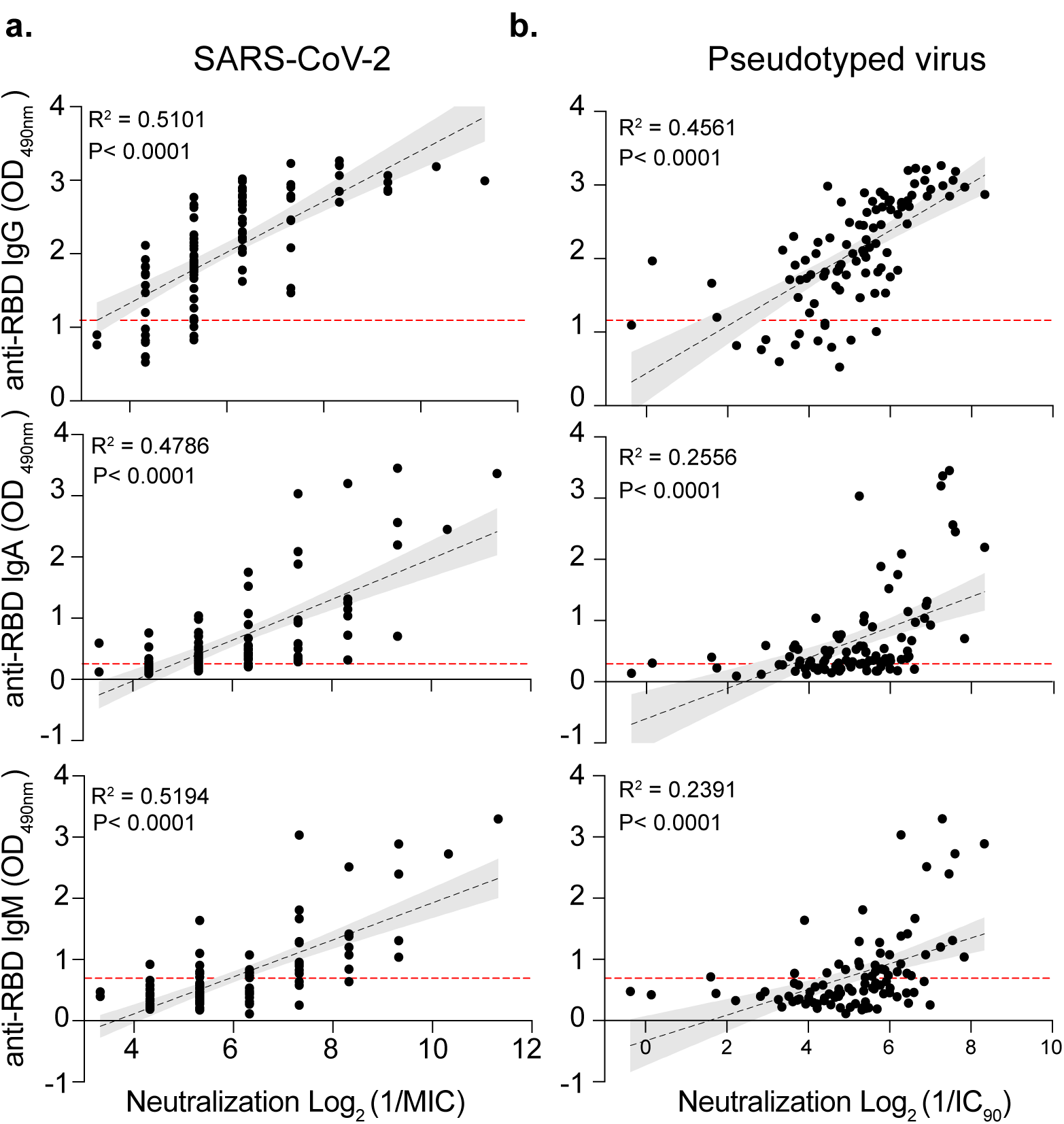
Correlation of anti-RBD antibody isotypes with viral neutralization. Anti-RBD ELISA correlation for IgG (top), IgA (Middle) and IgM (Bottom) with viral neutralization using authentic SARS-CoV-2 **(a)** or Pseudotyped virus **(b)**. Correlation and linear regression analyses were performed using GraphPad Prism 8. P values were calculated using a two-sided F-test. The red dashed lines indicate the threshold (anti-RBD ELISA for -IgG is OD490nm = 1.091; −IgM is OD490nm = 0.256 and −IgA is OD490nm = 0.694) for each ELISA.

### Robust SARS-CoV-2 neutralization associates with high-titer, multi-isotype antibody responses

Next, we delineated what separates our highest neutralizers from the remainder of the cohort using the data collected on antigen-specific antibody isotype titers. To do so, we employed a holistic approach and compared the IgG, IgM and IgA titers for both anti-Spike RBD and anti-N antibodies against neutralizations level (Fig. 4a and b). This revealed that the highest overall neutralizers (MIC higher or equal to 1/640) had higher titers of IgG, IgM and IgA raised against both Spike RBD and N (Fig. 4a and b, red bar). The spike protein is the major determinant for virus neutralization and resides on the outside of the viral particle, exposed to the immune recognition. Consequently, we found stronger correlations between Spike-RBD isotypes compared with N isotypes, when comparing each individual’s IgG, IgM or IgA (anti-Spike RBD and N) against each other (Fig. 4c and d). However, in most cases the high neutralizers (red circles) had the highest antibody titers for all isotypes (Fig. 4c and 4d). These data suggest that mounting a robust antibody response, consisting of diverse isotypes, leads to efficient neutralization.

**Figure 4.**
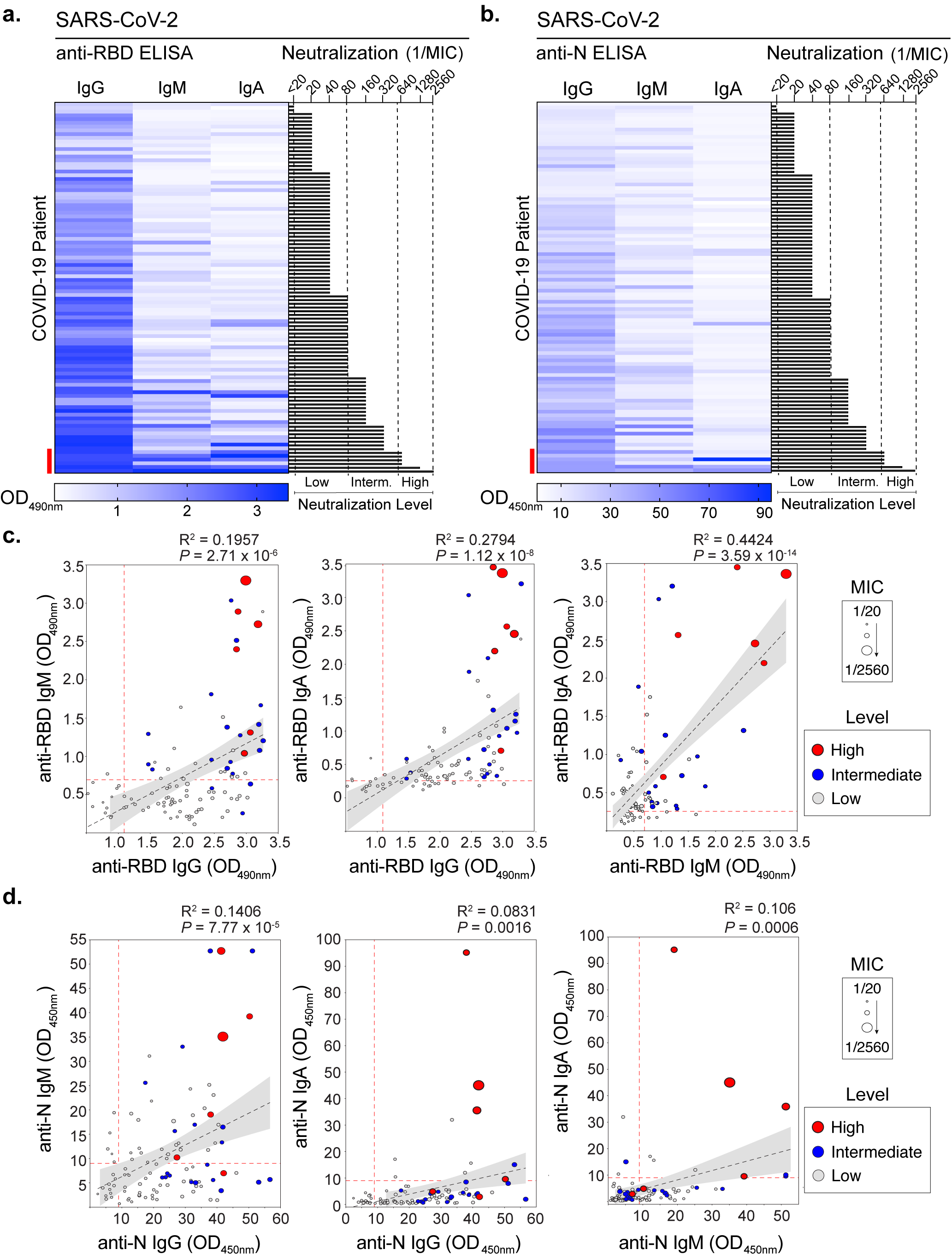
Identification of serological signatures for neutralization. Heatmap of anti-RBD **(a)** and anti-N **(b)** antibody isotype ELISA titers and corresponding authentic SARS-CoV-2 neutralization. Serological data from the 101 COVID-19 patients was ranked to high neutralization. Sera of convalescent patients was defined as low (dil:1/10 to 1/80), intermediate (dil:1/81 to 1/639) and high (dil> 1/640). Red bar indicates those COVID-19 patients with high neutralizer antibodies. c. Correlation analysis of anti-RBD IgM vs IgG (left panel), anti-RBD IgA vs IgG (middle panel) and anti-RBD IgA vs IgM (right panel). d. Correlation analysis of anti-N IgM vs panel), anti-N IgA vs IgG (middle panel) and anti-N IgA vs IgM (right panel). For (c) and (d) correlation and linear regression analyses were performed using the linear model function in R (lm). P values were calculated using a two-sided F-test. The size of the dots indicates the MIC and the color of the dots indicates the neutralization category: High (red dots), Intermediate (blue) and Low (gray) as determined using authentic SARS-CoV-2 neutralization (see Fig. 2 legend).

### Recovered individuals have multiple combinations of anti-SARS-CoV-2 serum antibodies

Finally, given that having multiple isotypes associate with higher neutralization, we were interested in understanding the anti-RBD and N antibody signatures present in individuals and how they correlated with neutralization. Therefore, we classified whether the presence or absence of certain antigen-specific antibody isotypes (based on ELISA cutoffs) related to neutralization level. The 101 COVID-19 patients comprised 21 different antibody combinations, or “clusters” (Fig. 5a), ranging from being positive for all six antibodies (N/RBD IgG, IgA, IgM) to three individuals who did not have antibodies to neither N or RBD (Fig. 5a and b, **Cluster U).** We found that 4 out of the 6 healthcare workers with high neutralizing titers (MIC = 1/640 – 1/2560) had sera that tested positive for all three isotypes (IgA, IgG, and IgM) responsive against both Spike RBD and N (Fig. 5b, **Cluster A)**. However, two individuals with high neutralizing titers (#61 and 67) contained a combination of isotypes that were shared with both medium and low neutralizers (Fig. 5b, **Clusters C and D)**. Interestingly, Cluster B, which only lacks IgM against the RBD does not have high neutralization compared with Clusters A, C, and D, suggesting that specific antibody combinations and/or titers may be necessary for maximum neutralization. Finally, the majority of individuals had at least IgG and IgA directed against the RBD accompanied by IgG to N (Fig. 5b). Taken together, these results suggest that having IgG alone may not be enough for efficient neutralization, while having multiple isotypes at high titers present in serum leads to more potent SARS-CoV-2 neutralization.

**Figure 5.**
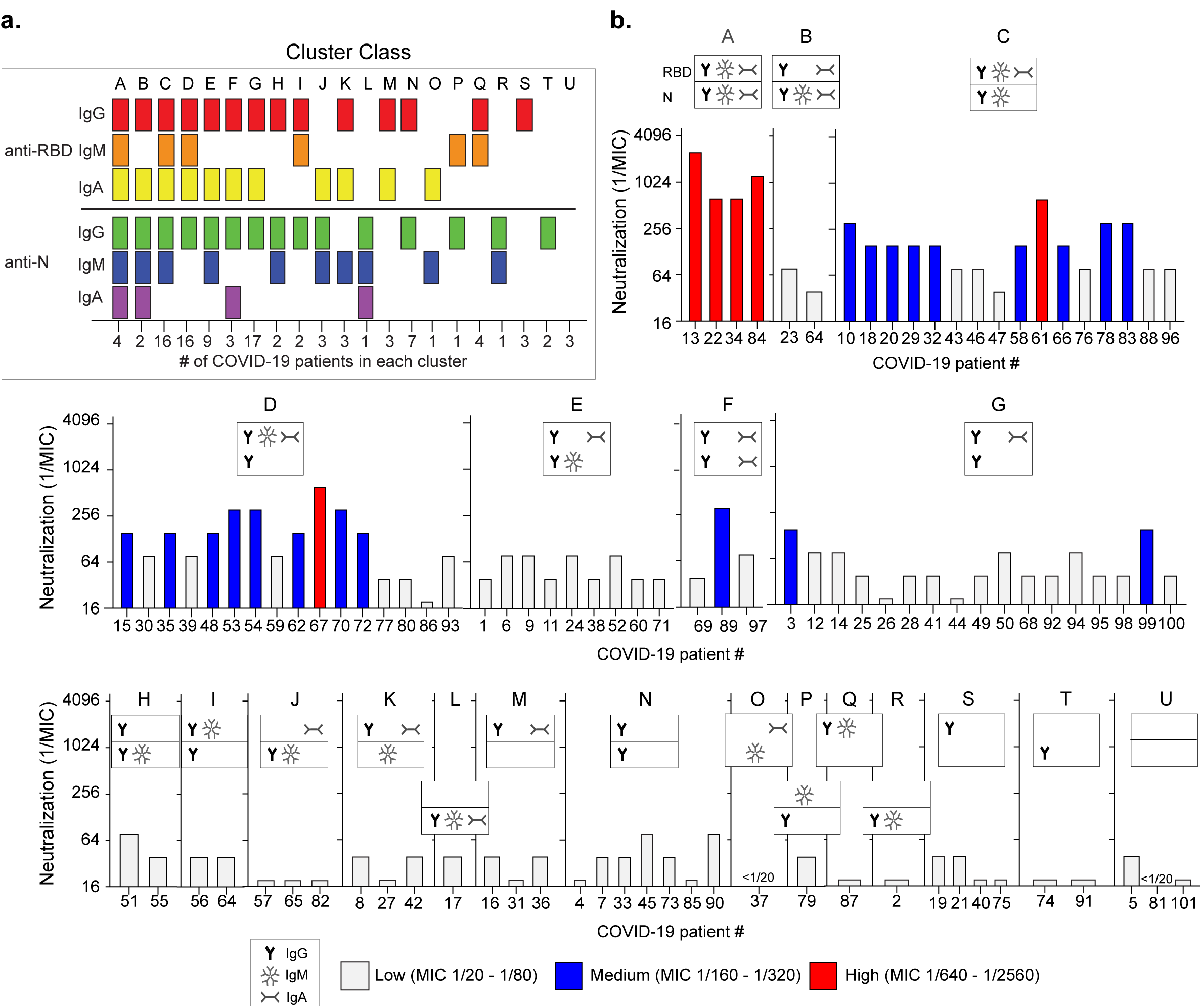
Clustering analysis of individual SARS-CoV-2 antibody response. **a.** Combination of antibody isotypes in individual patient sera defining different cluster classes. ELISA titers were categorized as positive (ELISA titers > cutoff) or negative (ELISA titer < cutoff) for each sample for each individual antibody isotype. Clusters were made based on the presence or absence of specific isotypes (Cluster A to U). The number of patients in each cluster is shown on the x axis. **b.** Neutralization levels shown for each individual antibody cluster.

## Discussion

Understanding the antibody response and potential immunity to SARS-CoV-2 is critical for global public health and the development of efficacious vaccines^1, 3, 26^. In this study, we performed a comprehensive analysis of the SARS-CoV-2-specific antibody response in 101 convalescent healthcare workers serum samples. We determined the neutralization capacity of the sera using both authentic virus and pseudotyped particles, quantified the titers of three antibody isotypes (IgG, IgM, and IgA) to both the spike receptor-binding domain and the nucleoprotein, and investigated the correlation of neutralization and antibody levels. We found, that the serum SARS-CoV-2 neutralizing antibody capacity was low for the majority of recovered individuals. In extreme cases, we detected no antibodies against SARS-CoV-2 Spike RBD or N in three individuals. These observations are in line with previously published studies and may suggest that those infected do not produce efficient neutralizing antibodies or neutralizing antibodies rapidly wane by the time samples are collected^16, 27, 28, 29^. Detailed longitudinal studies are beginning to emerge, showing that serum antibody levels are relatively stable or decrease over time30-32. In-line with our results these studies find a positive correlation between IgG titers and neutralization^31, 32^. Further characterization of neutralization at early and late timepoints is necessary to correlate antibody titer stability to viral protection. However, it is essential to keep in mind that, while neutralization titers can be quantified by reliable laboratory assays, we do not know the overall protective capacity of antibodies against SARS-CoV-2 in patients. Thus, those with low neutralizing capacities in the lab (MIC < 1/80) could be efficiently protected from SARS-CoV-2 reinfection. Detailed studies monitoring initial and possible reinfections along with the antibody response are crucial to understand immunity to SARS-CoV-2.

To further understand the composition of convalescent sera, we quantified the amounts of IgG, IgA, and IgM targeting the spike protein RBD and N. We found 21 different antibody combinations that did not directly correlate with days post symptom onset, yet did associate with antibody neutralization. Of these combinations, the majority of individuals fell into four distinct clusters comprised of different antigen-specific isotypes. It remains to be elucidated how these clusters are generated and why certain clusters elicit potent SARS-CoV-2 neutralization (Cluster A) and others do not (Cluster B). Since we use a targeted approach focusing on only two antigens and specific epitopes, it is possible that there are other anti-SARS-CoV-2 antibodies that bind viral particles and impact neutralization^33^.

Interestingly, we found high quantities of IgM present in some individuals from 32 to 50 days post symptom onset. These findings are intriguing since IgM antibodies are usually considered as a marker of a recent infection and their circulating titers are thought to decrease as class switching occurs to IgG and IgA. This observation is of importance for determining the relative time of infection. In this situation, looking at multiple isotypes including IgA as well as multiple antigens may be beneficial in determining the relative infection timeline. Along these lines, we also found that individuals having multiple antibodies isotypes to both RBD and N had the best neutralization. Moreover, we found that individuals who had IgG in combination with IgM and IgA against the RBD had the greatest neutralization capacity, suggesting that the generation of combinations of antibody isotypes against the RBD may provide the best neutralization. It has been shown that severity of disease can play an important role in the antibody response and it may be that in these individuals, disease is a driver for multiple isotypes and increased neutralization^15, 34^. Thus, understanding how enhanced disease burden leads to the generation of multiple neutralizing isotypes will be crucial for vaccine development.

These results, particularly the presence of individuals with low neutralizing antibodies are interesting as each individual in our cohort has recovered from SARS-CoV-2 infection. This observation suggests that either we missed the period of robust neutralizing antibody production in certain individuals or other mechanisms, such as potent SARS-CoV-2 specific T-cell responses^35, 36, 37–41^, provided effective viral clearance and immunity. Future studies coupling assessment of antibody and T-cell responses with key clinical information (age, sex, severity of symptoms) will be essential to fully understand this complex immune response. Moreover, these results also highlight the need to study the SARS-CoV-2 genetics as the virus encodes multiple undefined accessory proteins that could impact the immune response during infection. Additionally, longitudinal studies are necessary to fully understand the complex immune response to SARS-CoV-2 in the hopes to contain this virus and to prepare for the emergence of related viruses.

## Acknowledgements

We thank all the volunteers for participation in this study and the members of the NYU Vaccine Center and NYU Langone Health for obtaining these samples. We thank Drs. Chi Yun and Adriana Heguy for access to the Mantis instrument. We thank members of each lab for helpful comments and discussion on the study and manuscript.

---

**Extended Data Table 1:**
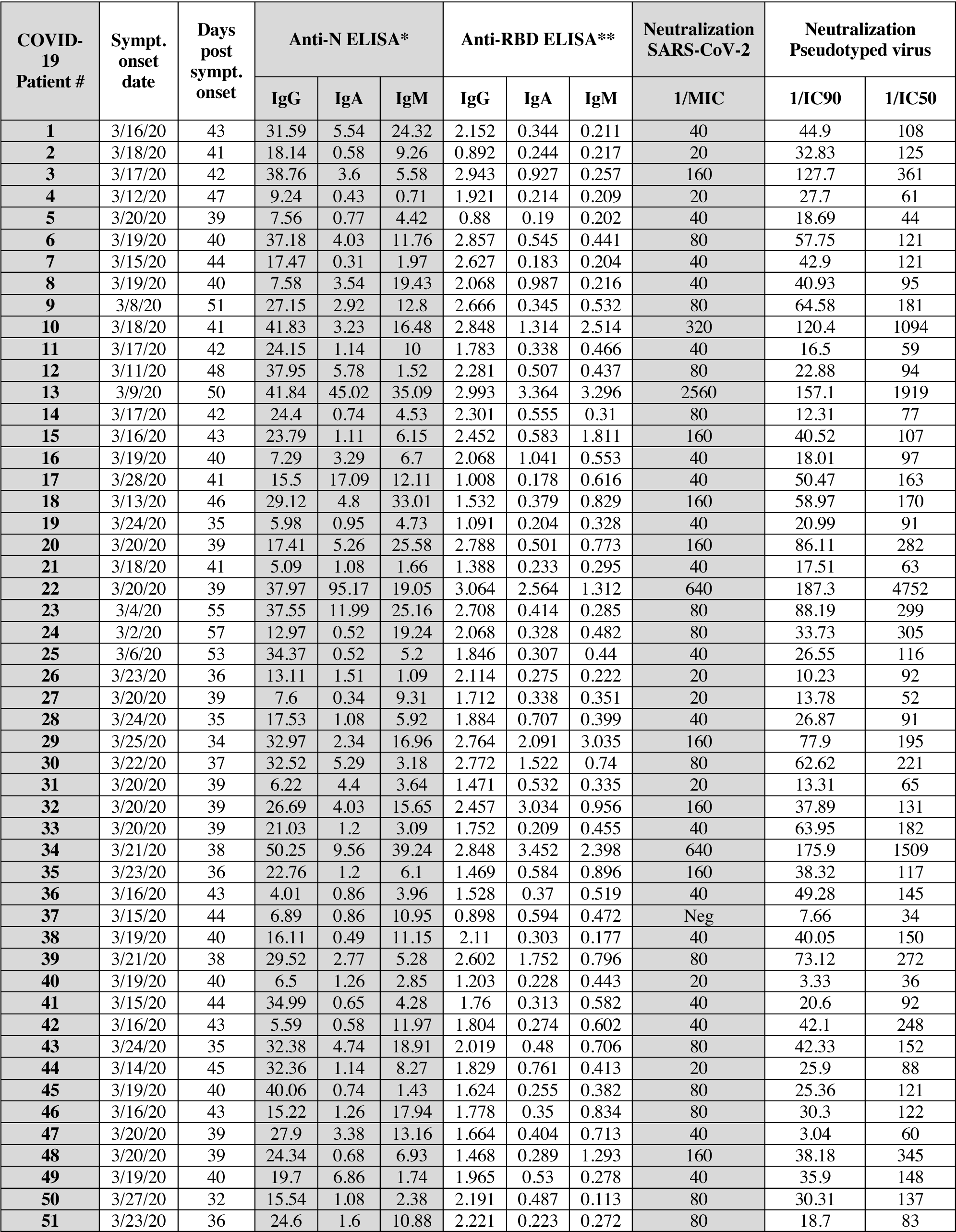
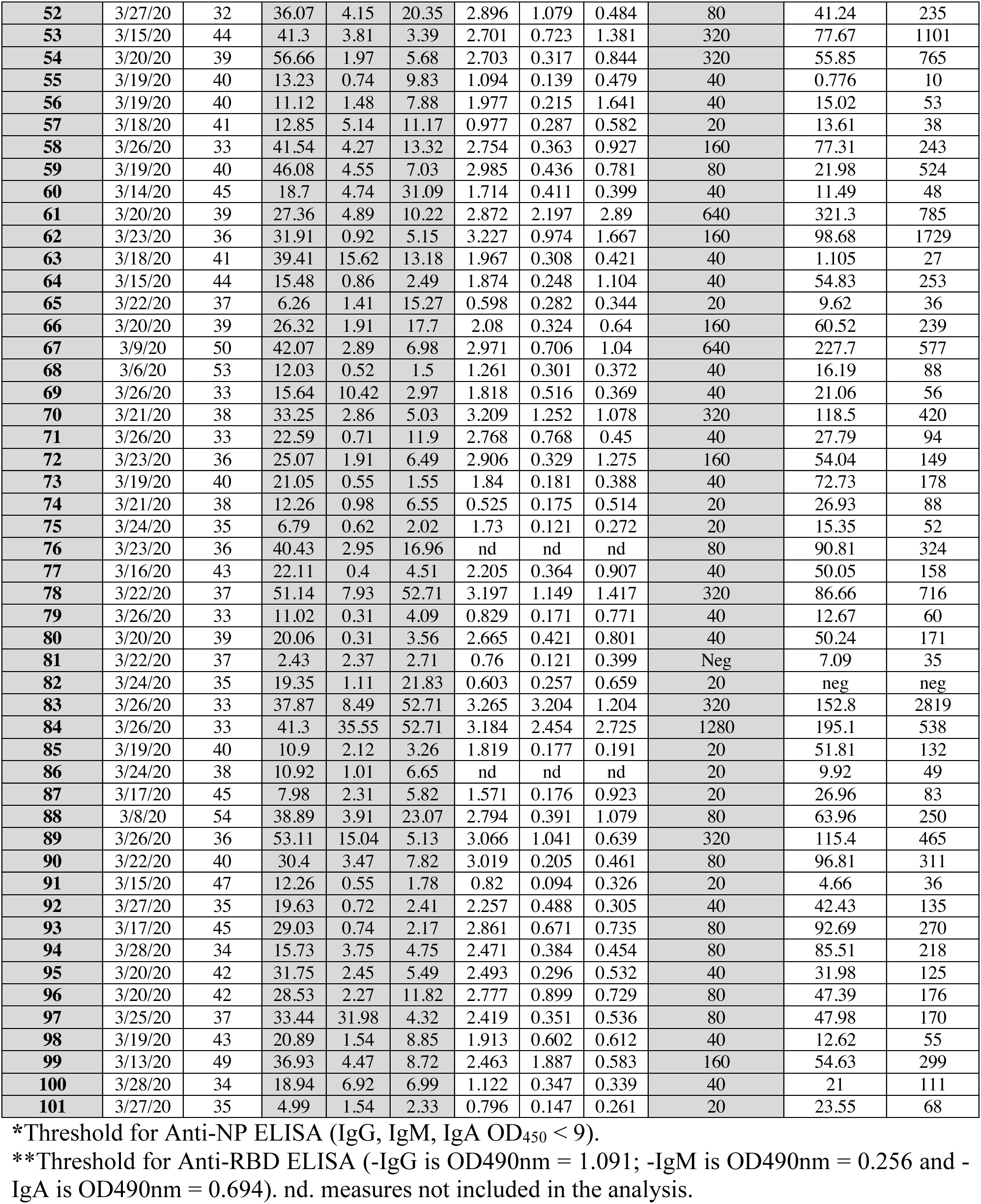
Extended Data SARS-CoV-2 positive NYU healthcare worker information and data used in this study.

**Extended data Figure 1.**
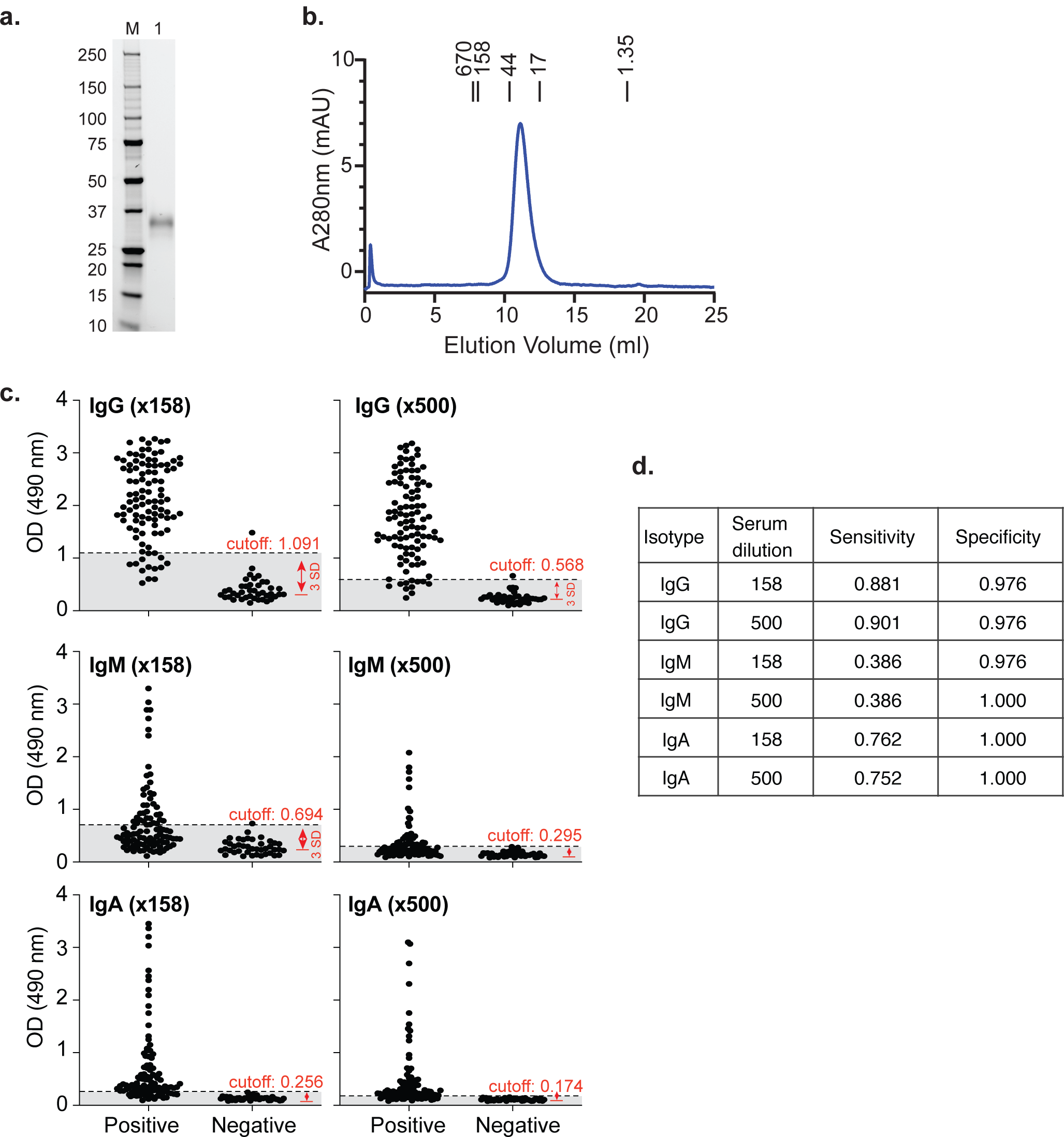
Production of SARS-CoV-2 Spike RBD antigen and characterization of anti-RBD ELISA. **a.** SDS-PAGE of purified RBD-His6-Avi-biotin using the Bio-Rad stain-free detection method. Lane M contains standards with its molecular weight in kDa. **b.** Purified RBD-His6-Avi-biotin size-exclusion chromatography on a Superdex 75 10/300 Increase column detected using absorbance at 280 nm. The elution positions of molecular weight standards are marked as bars with their molecular weights in kDa. **c.** Determination of ELISA thresholds. Serum from the 101 SARS-CoV-2 PCR positive were diluted 1/158 and 1/500 and compared with 43 SARS-CoV-2 PCR negative individuals. The cutoff values were defined as the mean plus three times the standard deviation (SD) of the negative control samples as shown by the red arrow. The dashed line indicates the position of each individual cutoff. Cutoff values are indicated in red. **d.** Table showing the sensitivity (% of true positive in positive) and specificity (% of true negative in negative) of each RBD antibody subtype and dilution.

**Extended data Figure 2.**
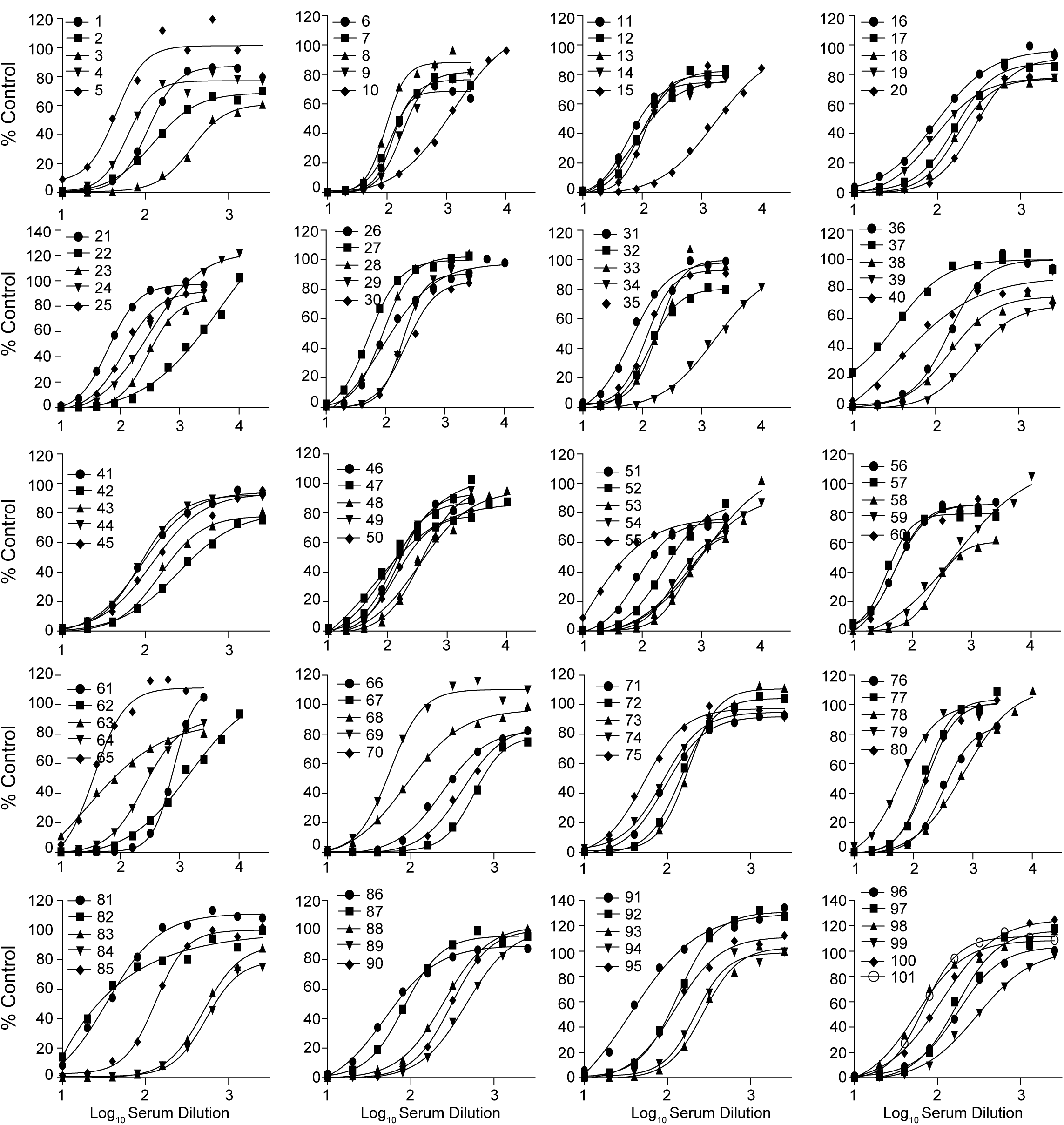
Individual sera neutralization of pseudotyped virus. Pseudotyped virus was 1:1 mixed with 2-fold dilutions of individual sera and incubated at room temperature for 30 minutes before infecting ACE2-293T cells. Relative infection was determined by luciferase expression 48 hrs post-incubation. Data is represented as the percentage of the untreated control. Data was fitted to a variable slope model log(serum dilution) versus response using Graph Pad prism.

**Extended data Figure 3.**
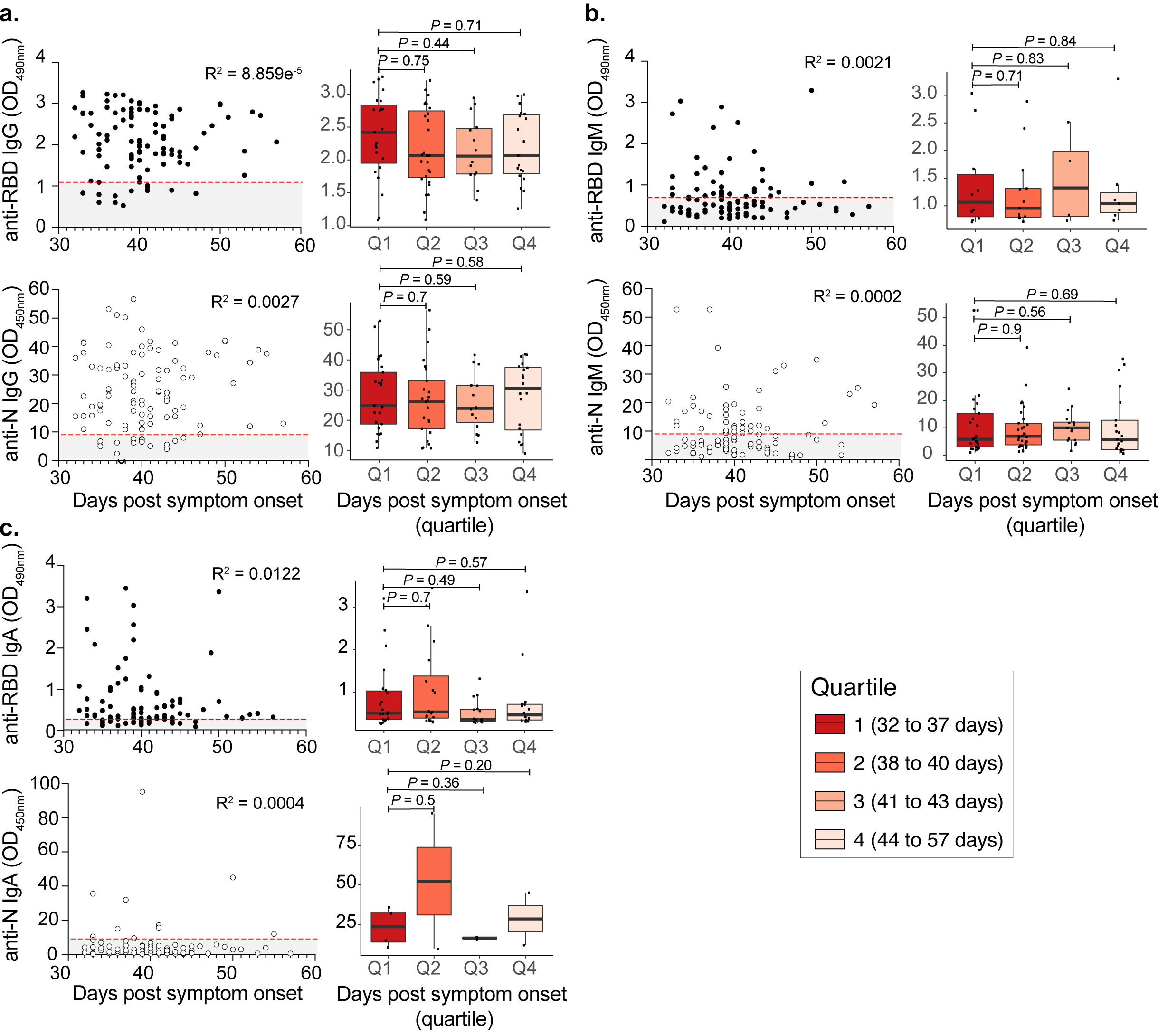
Correlations between isotype composition of SARS-CoV-2 convalescent serum and days post onset of symptoms. Correlations of anti-RBD (top) or anti-N (bottom) IgG **(a)**, IgM **(b)** and IgA **(c)** over days post onset of symptoms are shown. Quartiles were defined based on positive samples (data points > cutoff) for anti-RBD IgG. Each quartile has the following number of observations: anti-RBD IgG (n= 29, 30, 18, 22), anti-N IgG (n= 11, 13, 5, 8), anti-RBD IgM (n= 23, 20, 15, 17), anti-N IgM (n= 11, 12, 10, 8), anti-RBD IgA (n= 24, 28, 15, 20), anti-N IgA (n= 4, 2, 2, 2). P values were calculated using the Wilcoxon rank sum test. The red dashed lines indicate the threshold (anti-N ELISA for −IgG, −IgM or −IgA are OD450nm = 9; anti-RBD ELISA for −IgG is OD490nm = 1.091; −IgM is OD490nm = 0.256 and −IgA is OD490nm = 0.694) for each ELISA.

**Extended data Figure 4.**
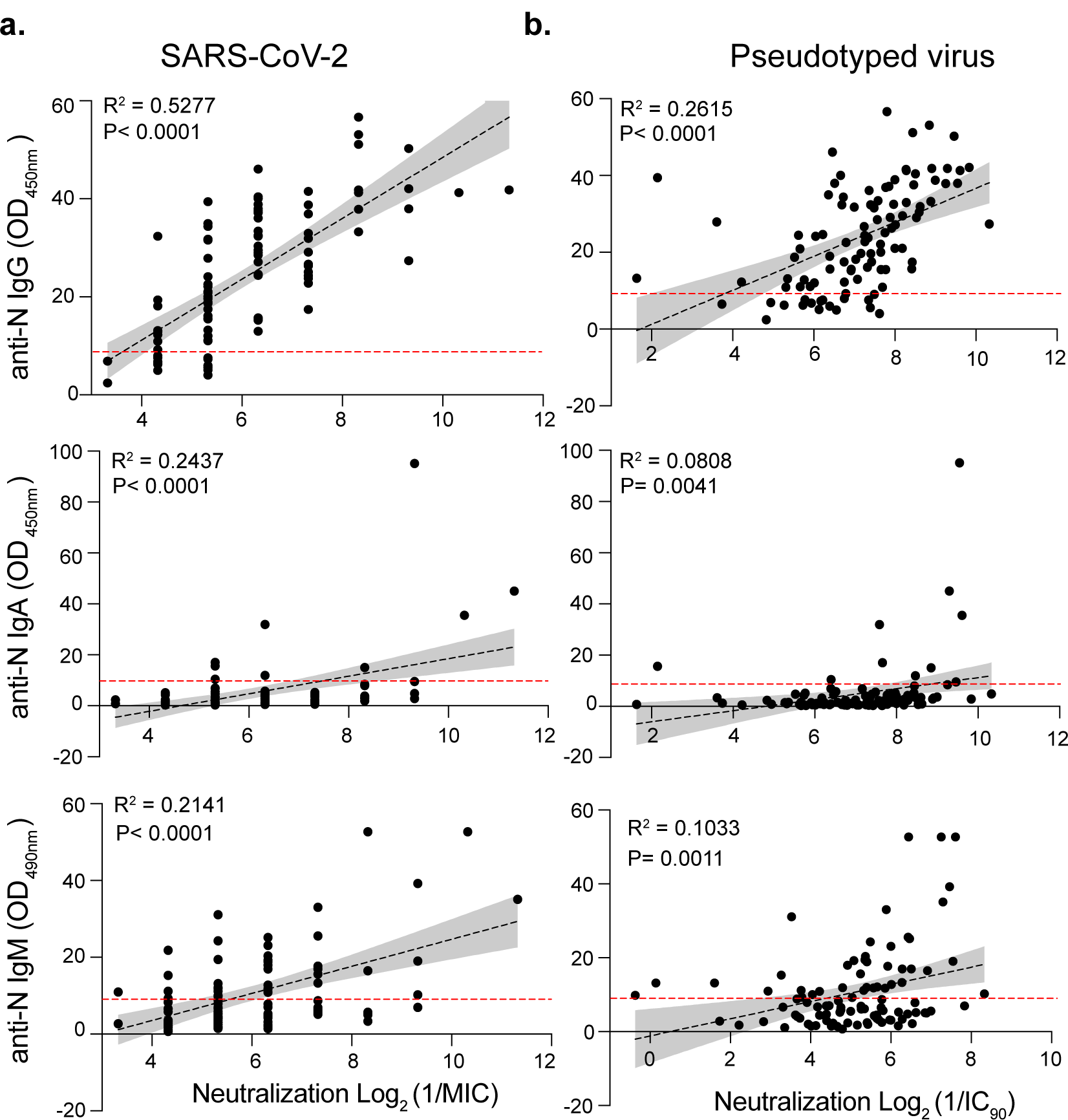
Correlation of anti-N antibody isotypes with viral neutralization. Anti-N ELISA correlation for IgG (top), IgA (Middle) and IgM (Bottom) with viral neutralization using authentic SARS-CoV-2 **(a)** or Pseudotyped virus **(b)**. Correlation and linear regression analyses were performed using GraphPad Prism 8. P values were calculated using a two-sided F-test. The red dashed lines indicate the threshold (anti-RBD ELISA for −IgG; −IgM and −IgA is OD450nm > 9) for each ELISA.

